# Machine learning as an effective method for identifying true SNPs in polyploid plants

**DOI:** 10.1101/274407

**Authors:** Walid Korani, Josh P. Clevenger, Ye Chu, Peggy Ozias-Akins

## Abstract

Single Nucleotide Polymorphisms (SNPs) have many advantages as molecular markers since they are ubiquitous and co-dominant. However, the discovery of true SNPs especially in polyploid species is difficult. Peanut is an allopolyploid, which has a very low rate of true SNP calling. A large set of true and false SNPs identified from the *Arachis* 58k Affymetrix array was leveraged to train machine learning models to select true SNPs straight from sequence data. These models achieved accuracy rates of above 80% using real peanut RNA-seq and whole genome shotgun (WGS) re-sequencing data, which is higher than previously reported for polyploids. A 48K SNP array, Axiom *Arachis*2, was designed using the approach which revealed 75% accuracy of calling SNPs from different tetraploid peanut genotypes. Using the method to simulate SNP variation in peanut, cotton, wheat, and strawberry, we show that models built with our parameter sets achieve above 98% accuracy in selecting true SNPs. Additionally, models built with simulated genotypes were able to select true SNPs at above 80% accuracy using real peanut data, demonstrating that our model can be used even if real data are not available to train the models. This work demonstrates an effective approach for calling highly reliable SNPs from polyploids using machine learning. A novel tool was developed for predicting true SNPs from sequence data, designated as SNP-ML (SNP-Machine Learning, pronounced “snip mill”), using the described models. SNP-ML additionally provides functionality to train new models not included in this study for customized use, designated SNP-MLer (SNP-Machine Learner, pronounced “snip miller”). SNP-ML is freely available for public use.

## Introduction

Single Nucleotide Polymorphisms (SNPs) are a major source of variation across plant genotypes. Therefore, the demand for discovery of a large number of SNPs increased after the advent of Next-Generation Sequencing (NGS). However, the extraction of true SNPs in polyploid organisms is challenging. Cultivated peanut is an allotetraploid, which poses an exceptional challenge for the discovery of true SNPs since the two parental diploid genomes (A and B) are very similar and the natural polymorphisms among peanut genotypes are very low [1,2].

Using Restriction-site-Associated DNA (RAD) sequencing, a large number of SNPs were identified in peanut diploid species; however, very few SNPs were discovered in cultivated peanuts [3]. Generally, the true SNP discovery in tetraploid peanut using NGS data is very low [4-6]. Sliding Window Extraction of Explicit Polymorphisms (SWEEP) was developed to improve the SNP calling by filtering out the polymorphisms between the two parental subgenomes [7]. However, SNP calling in tetraploid peanut requires additional improvement. An Affymetrix SNP array was designed using the SWEEP pipeline and showed promising genotyping results among cultivated peanuts [8]. The chip showed that SWEEP identified ∼40% true SNPs in tetraploid peanut genotypes. The array provided an unprecedented number of validated true and false SNP calls that can be leveraged with machine learning to increase the accuracy of selection of true SNPs straight from sequence data. The ability to have confidence in *in silico* SNP calls gives researchers access to all avenues of sequence-based genotyping methods.

Machine learning applies sets of different algorithms that facilitate pattern recognition and classification leading to prediction by creating models using existing data [9]. Machine learning algorithms are divided into two major classes; *i.e.* supervised and unsupervised. Supervised algorithms train previously well classified existing objects to predict the classes of new objects based on available features. Unsupervised algorithms cluster objects depending on their features without providing pre-defined classes. Both algorithms are used widely in different biological fields; *e.g.* coding region recognition, signal peptide prediction, biomarker identification, disease gene recognition, metabolic network detection, and protein-protein interaction [10-15]. For SNP calling, neural networks were used to differentiate between true SNPs and sequence errors and this method showed promising results for human SNPs [16]. In plants, neural networks also were used to classify called SNPs as true or false positives and the approach showed a positive prediction rate of 84.8% on the testing sets of soybean [17]. However, there has been little application of machine learning in polyploid organisms where the occurrence of more than one subgenome with high similarity to each other increases the complexity of read mapping and confounds the calling of true SNPs.

In this study, different supervised machine learners were used to improve the discovery of tetraploid peanut SNPs, utilizing the information of sequencing features and mapping data of the validated true- and false-positive SNP data sets extracted from analysis of the *Arachis* Affymetrix array. A new 48K SNP array was designed and validated based on the analysis of this method. Simulated SNP variant data from peanut, cotton, wheat, and strawberry also were used to extend the functionality of machine learning to other allopolyploids. Models trained with simulated data then were used to select SNPs from real peanut data with an accuracy exceeding 80%. This result has implications for using machine learning to select true SNPs in polyploid crops where no large validation sets are available. A tool was created, SNP-MLer (SNP-Machine Learner), which allows users to train models for use in selecting true SNPs from sequence data. The user can completely customize parameter sets used in training the models or default to the complete set used to train the peanut models. The models then can be implemented in SNP-ML (SNP-Machine Learning) to select true SNPs in new data sets.

## Materials and methods

### Data sets

The re-sequencing data set was created using 21 tetraploid *A. hypogaea* genotypes described in Clevenger et al. (2017) [8] and deposited publically at ncbi.nlm.nih.gov (Bio Project PRJNA340877 and Bio Samples SAMN05721179 to SAMN05721198). The RNA-seq data set has information from nine tetraploid peanut genotypes described in Clevenger et al. (2015, 2016a, 2016b) [18-20]. Validated true and false-positive SNP sets were based on testing the *Arachis* Affymetrix array with 384 peanut genotypes [8]. Mapping parameters were extracted from the vcf files used for the original design of the array. All positions of SNPs and surrounding sequence are based on the *A. duranensis* and *A. ipaensis* v1 pseudomolecules [https://peanutbase.org/, 1]

### Creating and testing a new Affymetrix array based on SNP-ML

A new affymttrix array was designed containing 28,218 SNPs which were extracted by SNP-ML using peanut real data re-sequencing model of neural network and tree bagger (S1 Table). The previously described 21 genotypes alongside 8 more genotypes and 103 minicore peanut lines [8] were assayed on the array and all 28,218 SNP-ML-derived markers were manually curated for polymorphism. A total of 21,112 markers were validated as polymorphic between genotypes (75%).

### Creating and testing the machine learning models

The data sets were prepared by R statistical software, *e.g.* extracting the attributes, randomly created training and testing sets and preparing fasta files for SNP flanking segments. Various toolboxes in MATLAB R2015b (the University of Georgia campus-wide site licensing agreement) were used for different purposes. Bioinformatics Toolbox was used for calculating the thermodynamic parameters, molecular weights and GC contents, Statistics and Machine Learning Toolbox was used for creating and testing the different models of supervised machine learning and Graphics functions were used for producing the bar plots and ROC (Receiver Operating Characteristic) graphs. The specific arguments of the different machine learning models are given in S2 Table.

### SNP-ML construction

We built paired (neural network and TB) specific trainer models for the two data types, WGS re-sequencing and RNA-seq. The models were built and stored in four files by a python script. In addition, three C++ classes were built, vcf.h, csv_write.h and csv.h, to process vcf and csv files. The SNP-ML main steps are illustrated in Fig 1. It uses C++ class vcf.h to extract the eight selected attributes from the input file, which is a vcf file of the output of SNP calling by mpileup of SAMtools, either directly or after primary filtration by SWEEP. The output is saved using the C++ class csv_write.h into a csv file, which is read by a python script to be applied to one pair of stored models (two files, one for neural network and the other for TB) depending on the data type. The two score sets are saved to a csv file, which is read by C++ class csv.h. The scores are filtered by passing only SNPs that have a value higher than the cutoff of neural network, which can be selected by the user (the default is 0.5), and occurred in the two score sets (shared SNPs in the output of the neural network and TB score file), in case the user selects that option. The scores are stored in csv files and the corresponding SNPs are stored in a vcf file.

**Fig 1:**
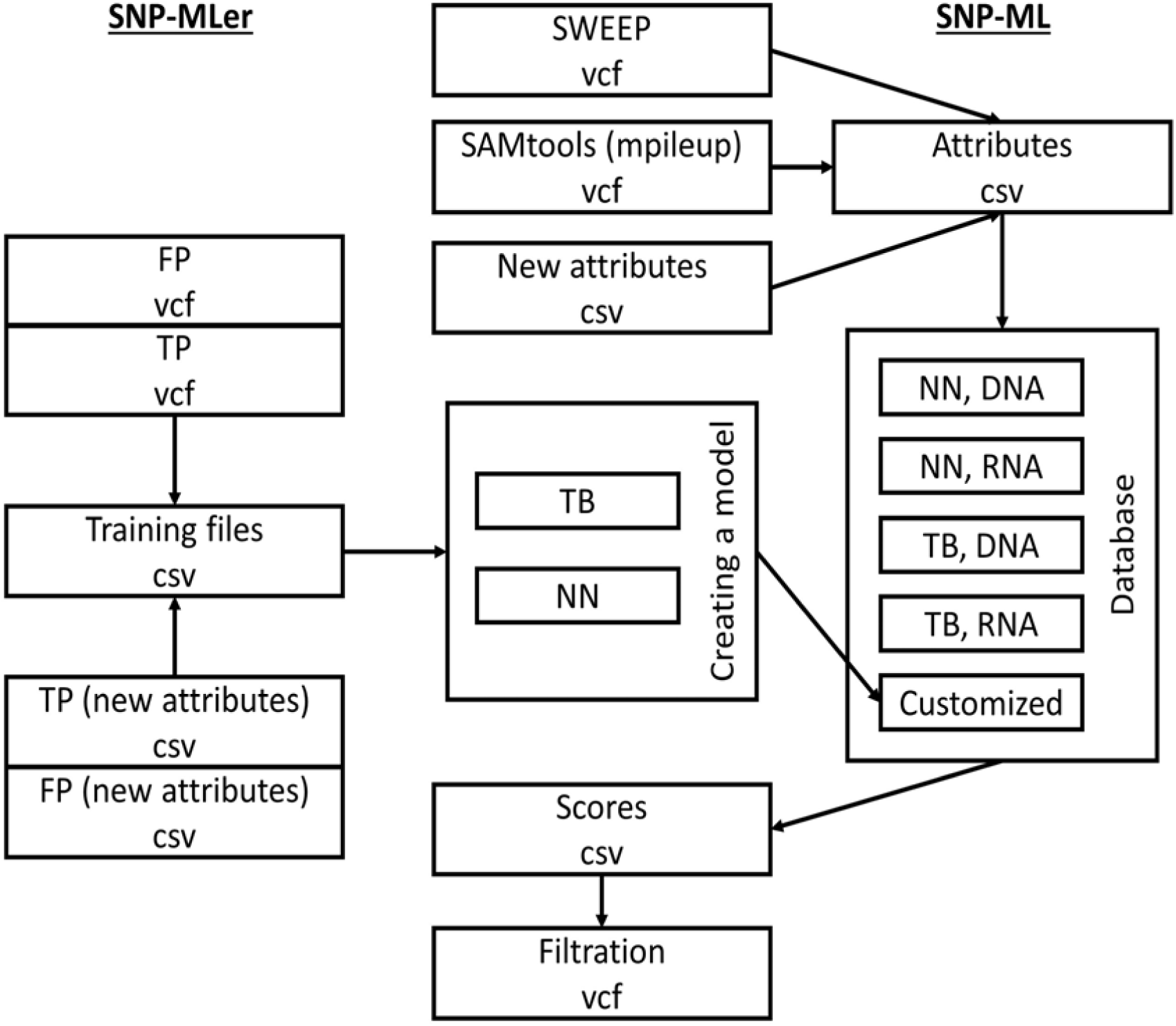
SNP-ML/SNP-MLer infrastructure. To extend the program applications, a second tool was designed, designated SNP-Mler (pronounced ‘snip miller’) to allow users to create predictors that are suitable for interested species/experimental conditions. SNP-MLer uses reading/writing approach as described above, it takes validated true-positive and false-positive vcf files as input and generates predictor model as outputs.

Both tools, SNP-ML and SNP-MLer allow the user to skip or select some of the eight attributes, and to apply new user defined attributes as csv files.

### SNP-ML requirements

The script was written by C++ and python 2.7.1 (S1 file). C++ was used for processing the data, input, output and filtering. The binary file was created by GCC 4.1.2 that was run on Red Hat 4.1.2-55 linux system. Python was used for creating the neural network and bagging machine learning models and applying the prediction using them. Different python packages were used for these purposes, i.e. numpy-1.11.0 (SciPy.org), scipy-0.17.1 (SciPy.org), pandas-0.18.1 (pandas.pydata.org), python-dateutil-2.0 (pypi.python.org), pytz-2016.4 (pypi.python.org), scikit-learn-0.17.1 (scikit-learn.org) and pyrenn 0.1 (pyrenn.readthedocs.io).

### Creating and testing models using simulated data

The pseudo molecule assembly AD1_BGI of cotton [http://www.cottongen.org/,21], the pseudomolecule assembly of the 3B chromosome of wheat [22], the contigs of TGACv1 wheat genome [https://plants.ensembl.org/index.html], the pseudomolecule assembly of *Fragaria vesca* Genome v1.1 [http://www.rosaceae.org/, 23], and the contigs of *F. nipponica* Genome v1.0 (FNI_r1.1), *F. nubicola* Genome v1.0 (FNU_r1.1) and *F. orientalis* Genome v1.0 (FOR_r1.1) [http://www.rosaceae.org/,24] were downloaded. 10,000 random loci were assigned in Chromosomes Aradu.A01, At_chr1, 3B and LG1, of peanut, cotton, wheat and strawberry, respectively. The loci were randomly mutated five times to form five synthetic genotypes using ART tool [25]. HiSeq 125 bp paired end sequences with different depths, 10x to 50x, were generated. The fastq produced files were mapped using BWA 0.7.10 [26] with default parameters to synthetic references as follows: a synthetic tetraploid reference containing Aradu.A01 and Araip.B01 chromosomes for peanut, a synthetic tetraploid reference containing At_chr1 and Dt_chr1 for cotton, a synthetic hexaploid reference containing 3B chromosome and the contigs of A and D genomes for wheat, and a synthetic octoploid reference containing LG1 chromosome and the contigs of FNI, FNU and FOR genomes for strawberry. SNPs were called using samtools mpileup 1.2 and bcftools 1.2.1 with default parameters without filtration. The SNP calling was carried out twice for every species. SNPs between two genotypes were called in the first instance and SNPs among the five genotypes were called in the second.

For each species, the SNPs located among the 10,000 loci were extracted in a separate vcf file, and considered to be True-positive (TP) SNPs. Any others identified by the program were extracted in another vcf file, and considered to be False-positive (FP) SNPs. Seventy percent of each one were randomly selected, and combined to be used as training sets, and the remaining 30% were used as testing sets for Neural Network models using Matlab R2015b (the University of Georgia campus-wide site licensing agreement).

### Testing simulated data against the real data

For peanut, 21 synthetic genotypes with 10X depth were generated and SNPs were called in four batches (three with five and one with six genotypes). The simulated data were used to train the model to mimic the conditions of the real data.

All sets of the TP and FP simulated data were used to train the models, to increase the strength, and the testing sets of the real data were re-applied to these simulated models. The generation of synthetic genotypes and carrying out the machine learning (training and testing) were applied as described above.

## Results and discussion

### Identification and evaluation of attributes for the model

A set of 18,057 validated true-positive SNPs and 26,050 false-positive SNPs were collected from the Axiom *Arachis* 58K SNP array [8]. These SNPs had been identified using SWEEP from 21 tetraploid peanut genotypes. The true-positive rate achieved was 40%, which was higher than previous efforts in peanut, but still inadequate. All of the mapping data in vcf form was available from the initial SNP calling, which provided the ability to test the hypothesis that machine learning would increase the accuracy of true SNP selection.

Seventy percent of the array-validated true- and false-positive SNPs (12,640 and 18,235, respectively) were randomly selected to train the machine learning model. Seventeen different attributes to be used in the model were calculated from sequences surrounding these SNPs (Table 1). These attributes were categorized into two groups, *i.e.* sequence and map features. The first machine learning approach used in biological applications was neural networks where it was used for recognizing the transcriptional start sites in *Escherichia coli* [9]. Since that time, it has become one of the most common machine learning approaches. In addition, neural networks have many advantages such as detection of all possible interactions between predictor variables, the ability to detect complex nonlinear relationships between independent and dependent variables, and applicability for different types of data sets [27]. Therefore, we used neural networks to build our first model and to select the most effective attributes.

**Table 1:**
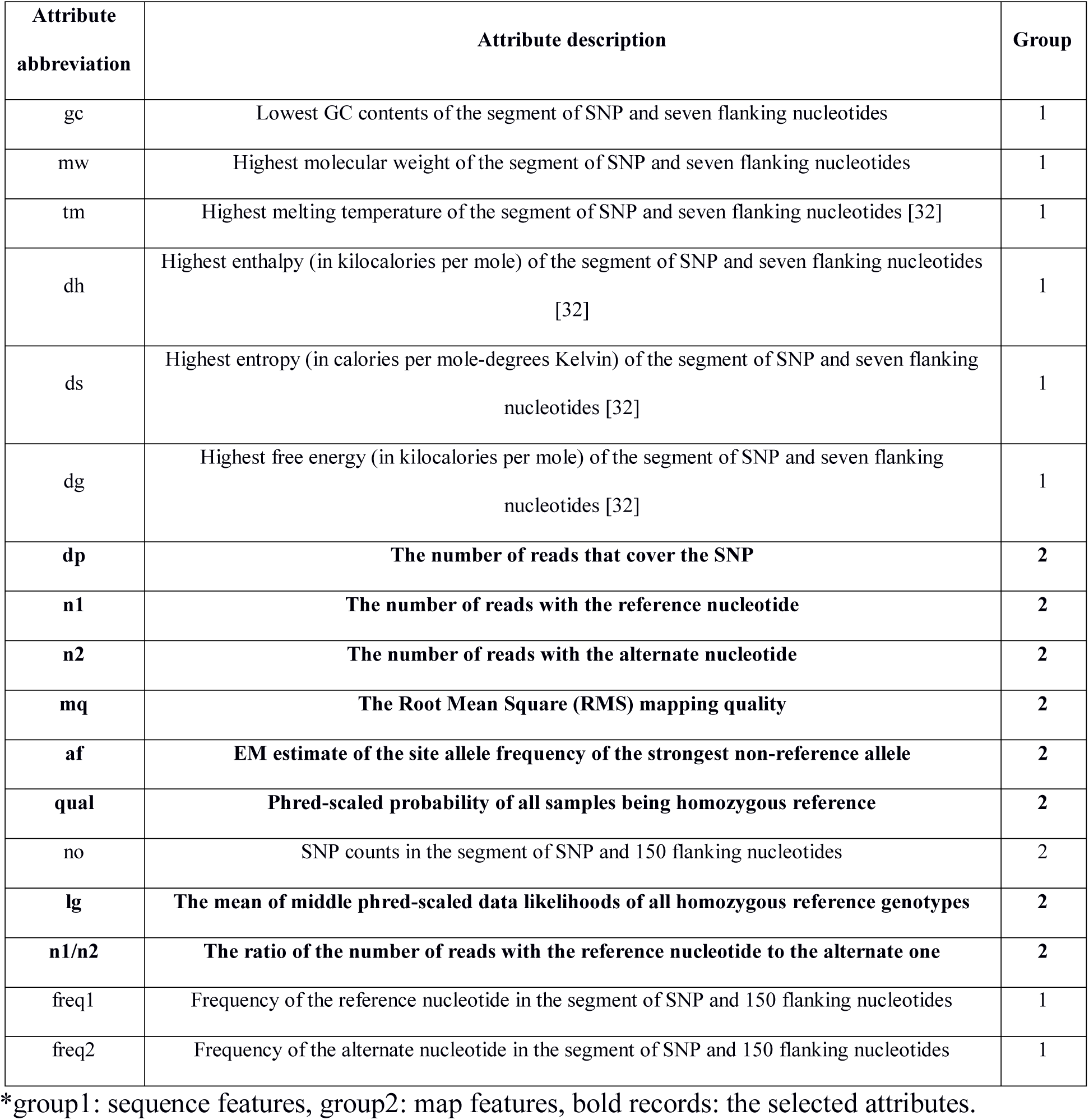
The attributes that were used for building the machine learning models.

Sequence features previously were used for genome wide *de novo* prediction purposes such as the prediction of coding regions, and to build a reliable neural network model for SNP calling in humans [16]. Thermodynamics of nucleic acids are important for diagnostic genetic markers for diseases, SNP sequencing on a genome-wide scale, designing PCR primers and creating probes for cloning and hybridization experiments [28]. Since thermodynamic parameters give indications for DNA molecule stability, they were used widely to predict the DNA secondary structure [29]. Therefore, we calculated the thermodynamic parameters deltaH, deltaS and deltaG for the SNP locations and flanking seven nucleotides (15 nucleotide segments) and incorporated the highest values from each pair of alternate SNP segments into the model. The higher value is associated with less stable states. Melting temperature (Tm) also was used in the same manner as it shares the primary components of deltaH and deltaS. Molecular weight was included since the change of a nucleotide affects the molecular weight of the DNA molecule. Lower GC contents decrease the stability of the DNA molecule. Therefore, we used the lower GC percentage of the two 15 nucleotide segments (the one with reference nucleotide versus the one with alternative nucleotide). In addition, frequency of the reference and alternate nucleotides in the sequences adjacent to the SNP location were calculated (for the seven nucleotides before and after the SNP location) and included in the model. We hypothesized that higher abundance of a particular nucleotide (reference or alternate nucleotide) would lower the probability of a true SNP.

The map features represent the quality of the mapping process and sequence data. Nine mapping parameters were selected to be used in the training model, namely quality features, *i.e.* mq (mapping quality) and qual (SNP quality); read abundance features, *i.e.* dp (depth of reads covering the SNP), af (minor allele frequency), n1 (reads with a reference base), n2 (reads with an alternate base) and n1/n2 (ratio of reference reads to alternate reads). In addition, a probability feature of homozygous reference genotypes (lg) was included. Some of these attributes, *i.e.* dp, n1, n2 and qual, were successfully used to create a neural network classifier for SNP calling for soybean [17]. Therefore, we assumed that these attributes and related features are good candidates for building a classifier in polyploids.

Twenty percent of the array-validated SNPs (3,611 true- and 5,210 false-positive SNPs) were used to test the model. Neural network models were applied to every one of the seventeen attributes independently and the relationship between false positives to false negatives was plotted for every model (Fig 2A). Interestingly, eight out of 17 attributes, all eight being map attributes, strongly affected the trainer (Fig 2A). These eight attributes were used for building one model, which showed a high reliability in classification of true- and false-positive SNPs (Fig 2B). The neural network score output of the testing data was applied to different neural network score cutoffs, from 0.1 to 0.9 by 0.1 intervals. The confusion matrices (predicted vs. actual) showed a gradual increase in the percentage of true negative (TN; false-positive SNP on the array and not called by SNP-ML) and decrease in the percentage of true positives (TP; true-positive SNP on the array and called by SNP-ML) as the cutoff increased (Fig 3). Increasing the cutoff over 0.5 dramatically decreased the percentage of TP SNPs, and also led to loss of a large number of valid SNPs (FN; true-positive SNP on the array but missed by SNP-ML). On the other hand, decreasing the cutoff below 0.5 increased the occurrence of a large number of false-positive SNPs (FP; false-positive SNP on the array and called by SNP-ML), an undesirable result. The cutoff of 0.5 showed a reasonable trade-off for recovery of the largest possible number of TP while minimizing FP and FN SNPs. These confusion matrices confirmed the efficiency of the eight selected attributes to build a reliable classifier.

**Fig 2:**
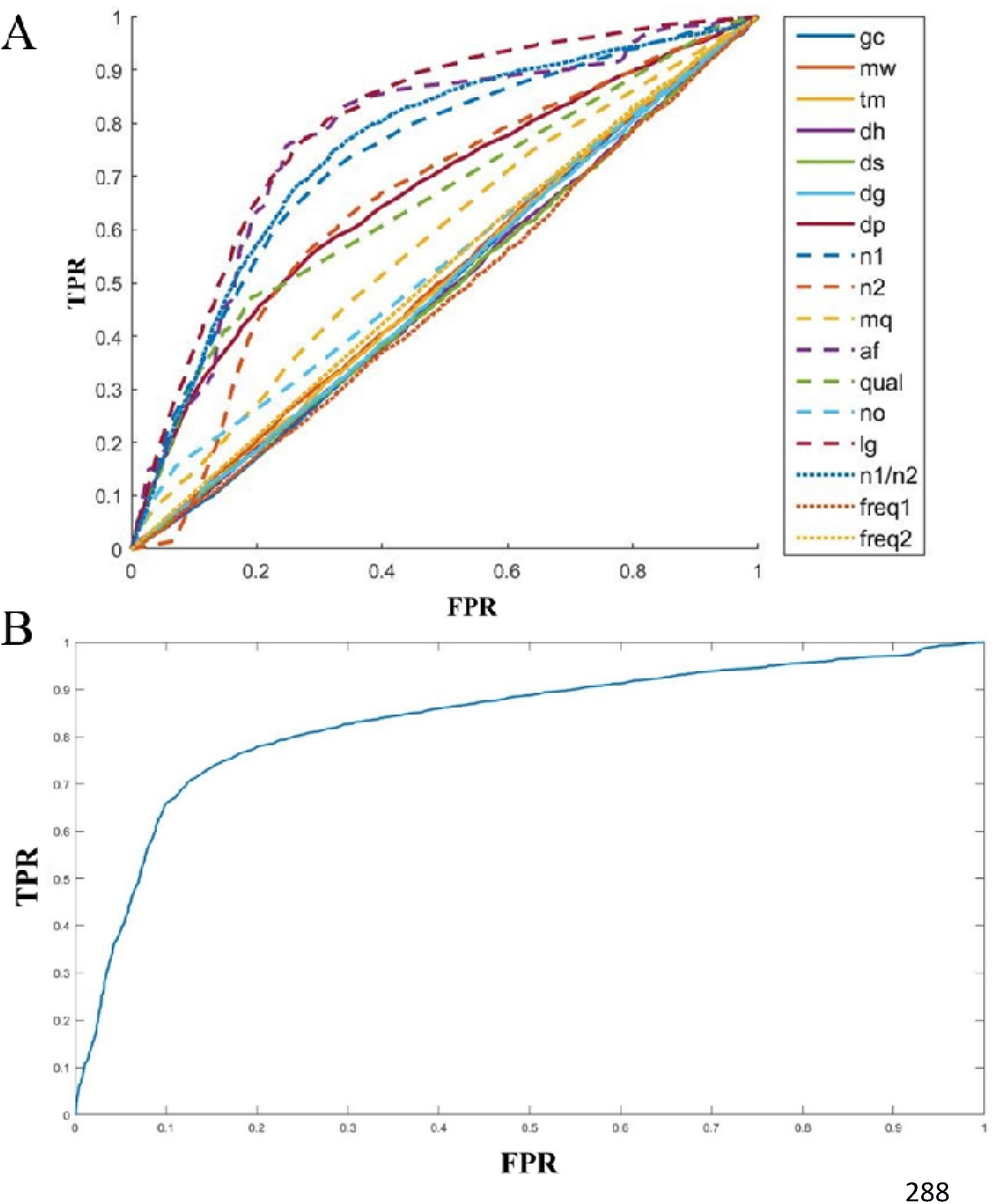
Receiver Operating Characteristic (ROC) curve of the attributes used in the neural network model trainer. A. Independent applications of the 17 attributes. B. The combined application of the selected eight attributes.

**Fig 3:**
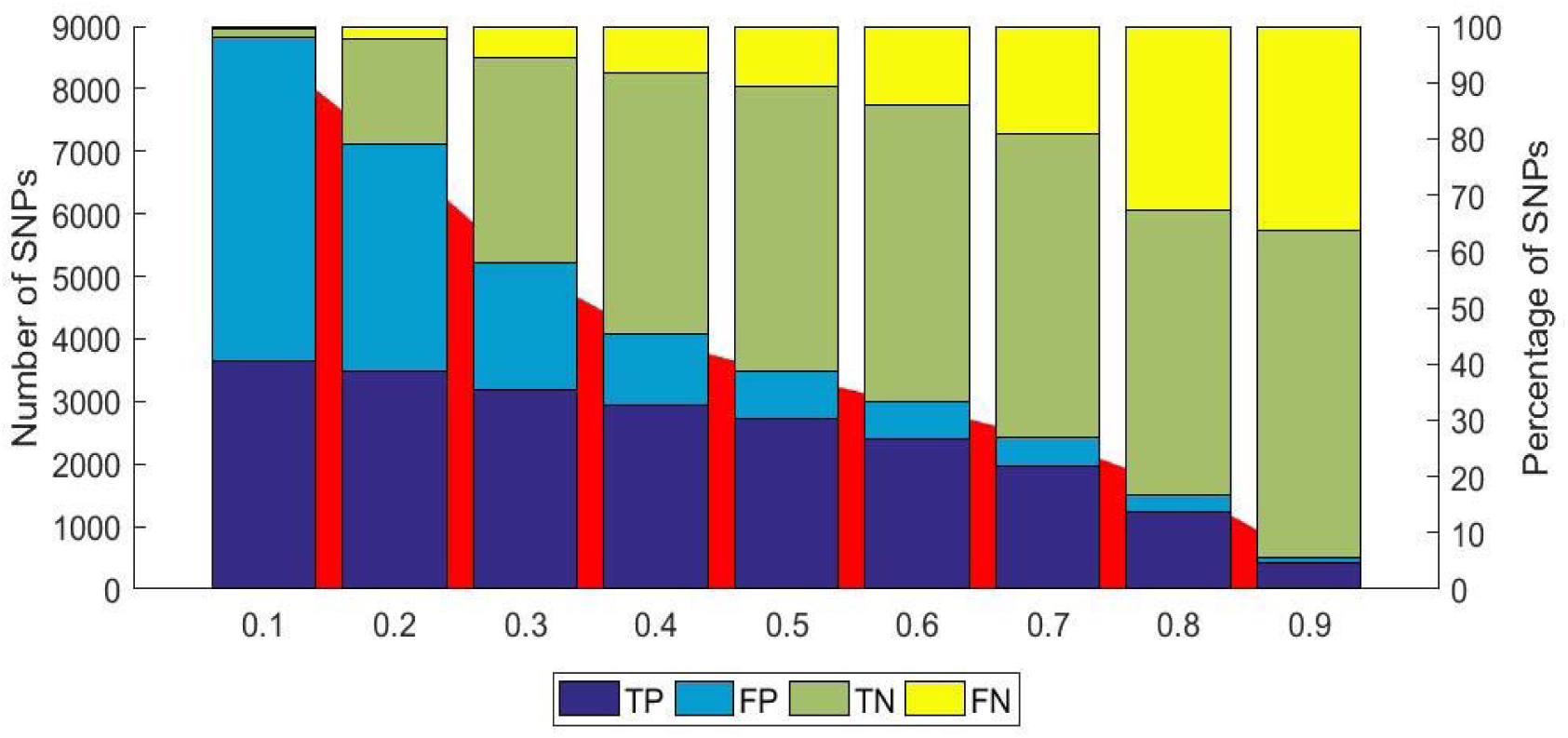
Bar plots representing the confusion matrices of the testing data using different cutoffs in neural network model, TP: True Positive (validated as a true SNP on the array and called by SNP-ML), FP: False Positive (not a true SNP according to array data but called by SNP-ML), TN: True Negative (not a true SNP according to array data and not called by SNP-ML), FN: False Negative (validated as a true SNP on the array and not called by SNP-ML). The red area shows the number of SNPs which are recognized by the model. The left Y scale present the number of SNPs within every class and the right Y scale presents the percentage SNPs of every class to the total SNPs.

### Comparison among different supervised machine learning models using the selected attributes

The training data set was used to build training models by applying different supervised algorithms, *i.e.* Logistic Regression (LR), Discriminant Analysis (DA), K-nearest Neighbors (KN), Naïve Bayes (NB), Decision Trees (DT) and TreeBagger (TB). The testing data set was applied for these trainers along with the neural network output of 0.5 cutoff (Fig 4). All models showed 60 to 80% true-positive rates relative to the number of SNPs extracted by a respective model, or between 25.0 to 33.4% of the total number of SNPs in the testing set. KN showed the highest false-positive rate and the neural network gave the lowest rate. Conversely, NB showed the lowest true-positive rate while TB produced the best rate. However, both TB and neural network showed the best trade-off between the two rates (Fig 5 and S3 Table). Therefore, we combined these two models to increase accuracy. TB was first described around 50 years after the first neural network approach was proposed [30]. It reduces the variance among observations and avoids overfitting, which are two limitations for neural network, thus it works as a complementary model to neural network to overcome its drawbacks.

**Fig 4:**
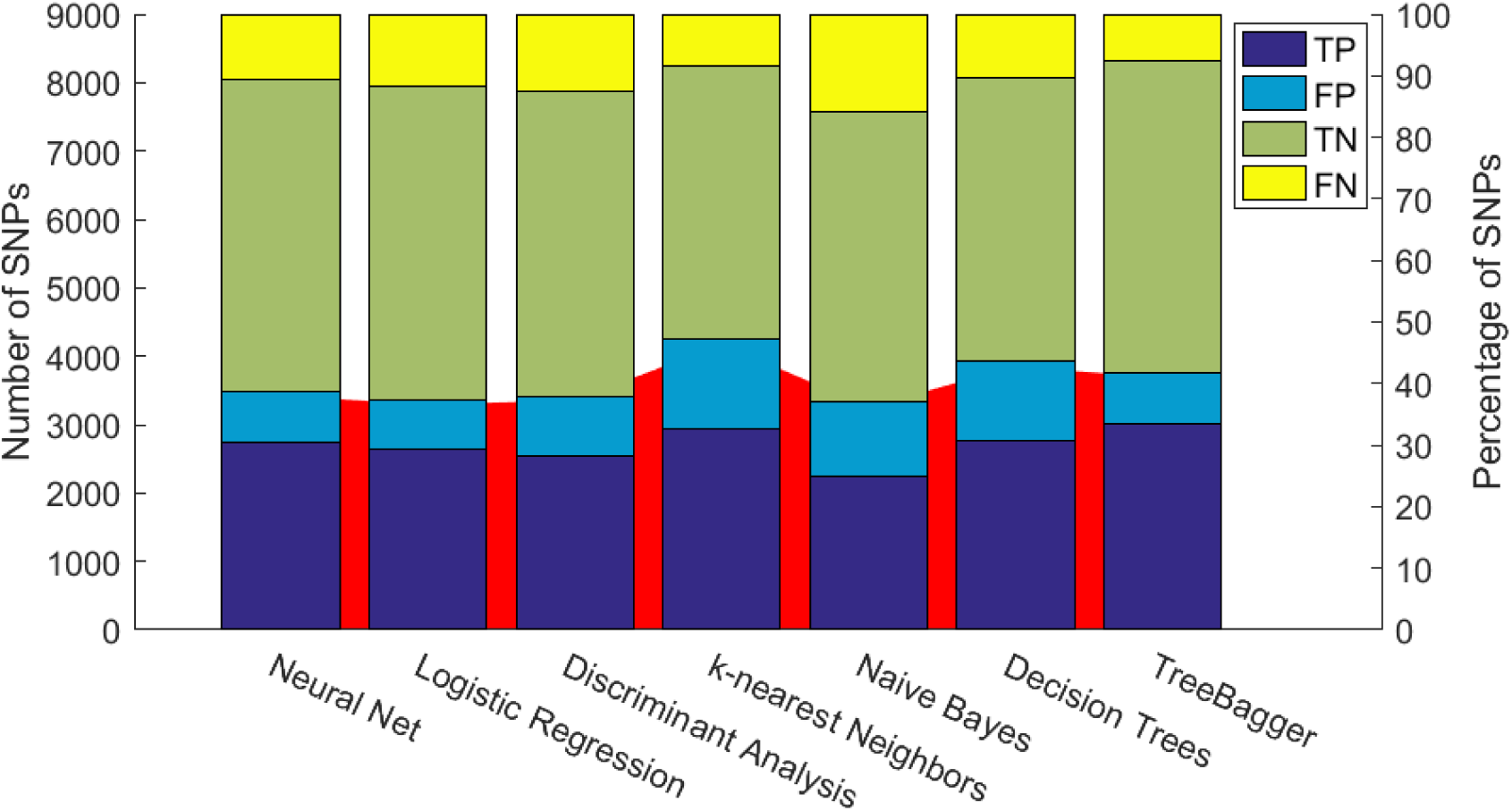
Bar plots represented the confusion matrices of the testing data using supervised machine learning algorithms, TP: True Positive, FP: False Positive, TN: True Negative, FN: False Negative. The red area shows the number of SNPs which are recognized by the model (positive total). The left Y scale presents the number of SNPs within every class and the right Y scale presents the percentage SNPs of every class to the total SNPs.

**Fig 5:**
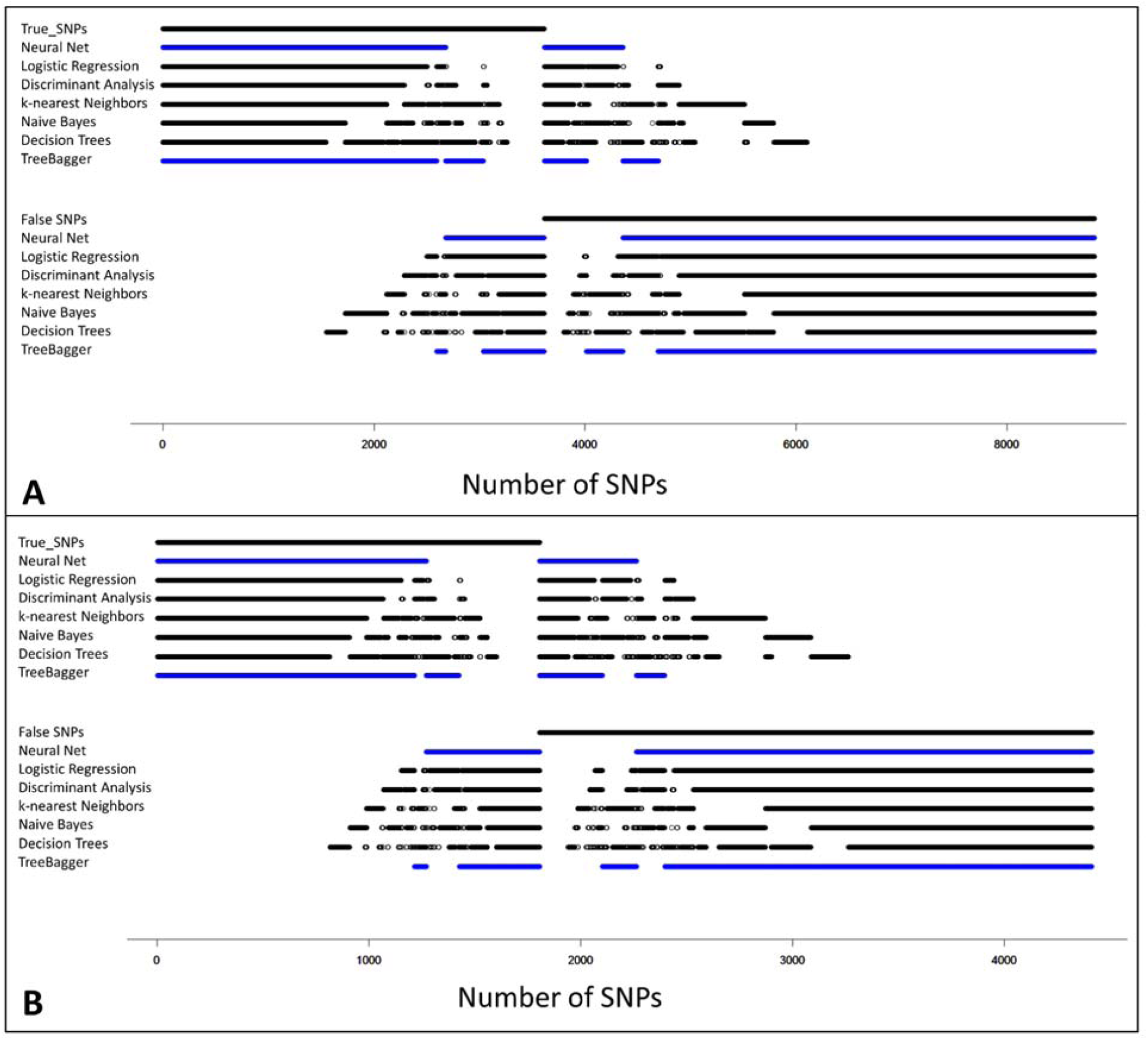
A dot plot of the trade-off combination between the different machine learning algorithms on the testing set (A) and validation set (B) , every dot shows a single true or false SNP (upper lines) and corresponding dots shows if this SNP was called using different machine learning algorithms.

To further test the model, the remaining 10% of the original data set, 1,806 validated true-positive and 2,605 false-positive SNPs, was used as a validation set. This data set was applied to the combined NN + TB model. A total of 1,510 SNPs was extracted by the model and 1,214 of those were true-positive SNPs. Therefore, the combined model efficiency increased to 80% versus 73% (1,271 out of 1,792) and 76% (1,369 out of 1,797) of using only neural network or TB, respectively. However, 33% of validated SNPs were lost through the prediction process using the combined model. The validation set of SNPs called using SWEEP and identified as true or false using the chip, is provided in S4 Table, along with detection state using only neural network, only TB, or the combined model.

### Model validation on Axiom *Arachis*2 48K SNP array

To validate the model for further real world analysis, 28,218 markers were selected to be included in a newly designed SNP array. A set of 133 tetraploid peanut genotypes and lines was genotyped on the chip. Polymorphisms were found in 21,112 SNPs among the tested genotypes, which revealed an accuracy of 75%. This represents the largest validation experiment to test a bioinformatics method developed to identify SNPs in polyploid species and provides the highest true positive validation rate reported in polyploids.

### Building models for RNA-seq

Unlike the re-sequencing data, RNA-seq provides data that measure gene expression and can produce a very high depth at specific loci [31]. The values of the attributes are different from the genomic re-sequencing data. For this reason, a specific model was built for RNA-seq data using sequence from nine tetraploid peanut genotypes. The analysis of this data set with SWEEP produced 3,525 SNP-chip overlapped SNPs, 2,143 true and 1,382 false SNPs.

Eighty percent of the array-validated SNPs were used for training the models, 1,714 true- and 1,104 false-positive SNPs, and the remaining 20% of SNPs were used as a testing set, 429 true- and 278 false-positive SNPs. Two models were built, *i.e.* neural network and TB, and the scored results were combined. The combined model extracted 371 SNPs (using the cutoff of 0.5 for neural network model). Of the SNPs extracted, 328 of them were true SNPs. The accuracy of true SNP discovery was raised to 88%. However, 101 SNPs were lost (∼24%).

### Application in other polyploids

In the absence of validation SNP sets for allotetraploid cotton (*Gossypium hirsutum*), allohexaploid wheat (*Triticum aestivum*), or allo-octoploid strawberry (*Fragaria x ananassa*), a simulation experiment was carried out to generate allelic variation. Genome sequence for each species was downloaded and five genotypes were simulated in one of the subgenomes while keeping the other subgenomes constant. The locations of true-positive SNPs thus were known due to the *in silico* mutation of the sequence and any other SNPs called by the program were considered false-positive. Because only one subgenome was mutated to derive the genotypes, all true SNPs were subgenome-specific. The true and false SNP calls were randomly categorized as training set (70%) and testing set (30%). The training set was used to train neural network models which were then used to select SNPs from the testing set. Simulations for all three species achieved accuracy of greater than 99% at five different sequence coverage depths (10X, 20X, 30X, 40X and 50X) (Table 2 and S5 Table). A peanut simulation was also included for comparison.

**Table 2:** The SNP-ML calling accuracy on different polyploid simulated data.

|  | Cotton | Wheat | Strawberry | Peanut |
| --- | --- | --- | --- | --- |
| Depth | True positive | True positive | True positive | True positive |
|  | % | % | % | % |
| **10X** | 100.00 | 99.64 | 99.72 | 99.85 |
| **20X** | 99.85 | 99.65 | 99.49 | 99.96 |
| **30X** | 100.00 | 99.88 | 99.73 | 99.96 |
| **40X** | 99.93 | 100.00 | 99.57 | 99.92 |
| **50X** | 99.96 | 99.88 | 98.15 | 99.96 |

### Application of simulation trained models on real data

Next, it was tested if models trained with simulated genotypes could achieve high accuracy in predicting true SNPs from real data, using the validation SNP sets available for peanut. Models that were trained in the simulations discussed above were used to select SNPs from the 21 genotypes of peanut (S6 Table). Each run of SNP-ML was performed three times to show variation between runs. For peanut, the models trained with simulated data were able to select true SNPs with accuracy on average of 78%. This result strongly suggests that this method can be used effectively in species where there are no large validation sets to train the models, but some reference sequence is available. This result, combined with the simulation results and results on real peanut data led us to construct a novel tool, SNP-ML, to carry out these analyses. The tool is designed to be highly flexible so that it can be used effectively in the broadest sense.

### SNP-MLer

All of the models discussed in this work are provided in the SNP-ML subdirectory “/db”. They include the peanut WGS and RNA-seq-trained models from real data and the models trained from simulated cotton, wheat, and strawberry data. The binary executable tool, SNP-MLer, will take two files as input, a vcf file containing true-positive SNP calls and a vcf file containing false-positive SNP calls. By default, SNP-MLer will train a neural network model using these sets of SNP calls and the eight parameters used in this work. The user has the ability with ‘-skip’ to not use one or more (up to seven) of the parameters if they wish or use ‘-custom’ to specify selected parameters in a comma-delimited sequence. Additionally, the user can use ‘-m’ to train a treebagging model as well. Most importantly, the user can add customized parameters to include in the model training by invoking ‘-addnew1’ and ‘-addnew2’. These options take csv files that include one or more new parameter lists for the true-positive SNP calls (-addnew1) and the false-positive calls (-addnew2). The user also needs to add the prefix name for the new model using ‘-o’.

### SNP-ML

If the user has trained new models using SNP-MLer or will use the models trained in this study, all models are located in the ‘\db’ folder for use with SNP-ML. SNP-ML is the tool that will take as input (-i) a vcf file of the SNP calls of interest. It is recommended to first use SWEEP to filter most of the false-positive SNP calls, but it is not required. The name of model to be used for SNP selection (-iM) should also be given as input to SNP-ML. The program contains currently two models, “peanut_DNA” for use with WGS data, and “peanut_RNA” for use with RNA-seq data. Any new models trained with SNP-MLer by the user will be included in this folder as well.

Users can submit any newly trained models to be included in new versions of SNP-ML by emailing the author. SNP-ML has similar options as SNP-MLer to skip (-skip) or customize (-custom) parameter sets for SNP prediction, and to invoke the treebagging model (-m) or add new parameters (-addnew; for custom trained models). An additional option (-c) allows the user to increase or decrease the stringency of true-positive selection from the default of 0.5. As discussed above and in Fig 3, increasing this cutoff will decrease false positives (decreasing selection of false SNPs) while increasing false negatives (limiting recovery of validated true SNPs) while decreasing the cutoff has the opposite effect.

The program is freely available for public use under MIT license and can be downloaded from https://github.com/w-korani/SNP-ML. A help file containing detailed information about using the program can be accessed by typing SNP-ML -h.

## Conclusions

We introduce a highly reliable method for calling SNPs for polypoid species using machine learning. To have a good classifier, the most effective attributes should be determined. Many attributes were tested and the best were selected for creating the model. In addition, different supervised machine learning algorithms were tested and the best ones for the data sets, neural network and bagging, were combined. We built and tested our method on peanut, an allotetraploid for which identifying true SNPs has been difficult. In addition, a 48K SNP array was designed using SNP-ML was created and showed high accuracy. The method was then used on simulated data from three other allopolyploids with different ploidy levels and achieved high accuracy. Most importantly, we showed that simulated data can be used to train models that achieve similar accuracy in selecting true SNPs using real data as do models trained with real data. The implication is that for species where there are no large validation sets available, our method can still be used to efficiently select true SNPs. With this important result in mind, SNP-MLer was developed; a tool that will train new neural network or treebagging models with user inputted data. Subsequently, SNP-ML can be used with newly trained models or included peanut models to select true SNPs for two different data set types, re-sequencing and RNA-seq. The flexibility and functionality of these tools allow the user a completely customizable experience, giving the ability to use the power of machine learning to researchers of all expertise levels.

## Acknowledgments

This work was supported by the Peanut Foundation, the Agriculture and Food Research Initiative competitive grant 2012-85117-19435 of the USDA National Institute of Food and Agriculture and the Feed the Future Innovation Lab for Collaborative Research on Peanut Productivity and Mycotoxin Control (Peanut and Mycotoxin Innovation Lab), supported by funding from the United States Agency for International Development (USAID).

## Availability of data and materials

SNP-ML, SNP-MLer and extendable database are freely available for public use under MIT license and can be downloaded from https://github.com/w-korani/SNP-ML

## Authors’ contributions

WK collected sequence attributes, applied the models, programmed SNP-ML/SNP-MLer and drafted the manuscript. JC collected map attributes, designed SNP array, edited and revised the manuscript. CY collected the validation data of the new SNP array. PO conceived and supervised the project, secured funding, and revised and submitted the manuscript.

## Supporting information

S1 Table: Affymetrix SNP array structure.

S2 Table: The specific arguments of the different machine learning models.

S3 Table: The efficiency of the different machine learning models.

S4 Table: The validation set of SNPs called using SWEEP, neural network, TB, and the combined model.

S5 Table: The efficiency of SNP-ML SNP calling for simulated data of cotton, wheat, strawberry and peanut.

S6 Table: The efficiency of using simulated data model to call SNPs from real data in peanut.

S1 File: The source code of C++ and python of SNP-ML (SNP-ML_source_code.zip).

